# ARIC: Accurate and robust inference of cell type proportions from bulk gene expression or DNA methylation data

**DOI:** 10.1101/2021.04.02.438149

**Authors:** Wei Zhang, Hanwen Xu, Rong Qiao, Bixi Zhong, Xianglin Zhang, Jin Gu, Xuegong Zhang, Lei Wei, Xiaowo Wang

## Abstract

Quantifying the cell proportions, especially for rare cell types in some scenarios, is of great value to track signals related to certain phenotypes or diseases. Although some methods have been pro-posed to infer cell proportions from multi-component bulk data, they are substantially less effective for estimating rare cell type proportions since they are highly sensitive against feature outliers and collinearity. Here we proposed a new deconvolution algorithm named ARIC to estimate cell type proportions from bulk gene expression or DNA methylation data. ARIC utilizes a novel two-step marker selection strategy, including component-wise condition number-based feature collinearity elimination and adaptive outlier markers removal. This strategy can systematically obtain effective markers that ensure a robust and precise weighted υ-support vector regression-based proportion prediction. We showed that ARIC can estimate fractions accurately in both DNA methylation and gene expression data from different experiments. Taken together, ARIC is a promising tool to solve the deconvolution problem of bulk data where rare components are of vital importance.

## Introduction

High-throughput biological technologies, such as microarrays, RNA-seq and whole-genome bisulfite sequencing, provide us informative approaches to investigate various biological samples collected from laboratories or clinical trials [1, 2]. However, plenty of these biological samples are complex mixtures of many different cell types without knowing their accurate proportions. Meanwhile, almost all physiological and pathological processes in multicellular organisms involve multiple cell types, each playing its particular roles [3]. Therefore, estimating the proportions of all or some specific cell types from bulk high-throughput data helps to understand the mechanism of biological processes and decouple signals related to certain phenotypes or diseases. For instance, the proportion of white blood cells can indicate the severity of immunological rejection after transplanting kidneys [4]. Quantifying the density of tumor infiltrating lymphocytes (TILs) like CD3^+^ and CD8^+^ T cells helps to predict patient survival in kinds of cancers [5-9]. Estimating placental signals in plasma cell-free DNA (cfDNA) of pregnant women can characterize the development of fetuses [10]. Detecting tumor-derived DNA fragments from plasma cfDNA can help us reveal the origin and development of cancers [11].

The fractions of minority cell types are of special interest in some deconvolution tasks. For example, the fraction of tumor-derived cfDNA in total plasma cfDNA, which is ultra-low for patients with early-stage cancer, is of vital importance for tumor detection [12]. TILs, which are promising biomarkers for clinical outcome prediction, also exhibit low fractions in many cancer tissues [5, 7, 13, 14]. Precise estimation of the fractions of certain cell types, especially rare cell types for some scenarios, could benefit a lot to discover signatures related to certain phenotypes or diseases.

Currently, many cell-type-specific omics data such as DNA methylation and gene expression profiles have been produced and can be accessed from various resources, like The Cancer Genome Atlas (TCGA) program [15], Encyclopedia of DNA Elements (ENCODE) project [16] and Gene Expression Omnibus (GEO) [17]. These data help to extract unique features of specific cell types, laying the foundation for estimating their fractions from bulk data.

Several cell type deconvolution methods have been proposed recently. Many researchers modelled this problem as a mixing process of biological signals from each cell type and employed overdetermined linear equations to solve it [18-22]. For example, CIBERSORT held bulk gene expression data as a linear mixture of different cell types [18]. The stability of the signature matrix was measured through the 2-norm condition number and support vector regression (SVR) was adopted to resolve the proportion of each cell type successfully [18]. Moss *et al*. proposed a linear model based on external references for DNA methylation data which successfully estimated the contribution of tissues to cfDNA [20]. Reference-free deconvolution methods such as non-negative matrix factorization (NMF) also perform well in some situations [23, 24]. Besides, some other studies modelled the mixing process with probabilistic models. For instance, PERT regarded the mixture of gene expression as a result of a sampling process and computed cell type proportions by the non-negative maximum likelihood model [25]. CancerLocator and CancerDetector regarded DNA methylation as a beta distribution to predict the fraction of tumor-derived reads in cfDNA sequencing data [26, 27].

It is rather remarkable that the above-mentioned studies did not pay sufficient consideration to rare cell types. Rare cell types have relatively low proportions, therefore, deconvolution for these cell types are more prone to be biased by collinearity. In addition, these studies did not provide efficient methods to avoid the impact of outliers during the selection of cell-type specific markers, preventing themselves from accurate deconvolution [19, 28], especially for rare cell types.

Here, we presented ARIC as a new approach for robust and accurate inference of rare cell type proportions from bulk gene expression or DNA methylation data. ARIC adopts a novel two-step feature selection strategy to ensure an accurate and robust detection for rare cell types. ARIC introduces the component-wise condition number into eliminating collinearity step to pay equal attentions for the relative errors of all components. Besides, ARIC contains an automatic step for adaptively removing the outliers in markers, ensuring the robustness of the algorithm against noises. Finally, ARIC employs a weighted υ-support vector regression (υ-SVR) to get component proportions. We evaluated ARIC in DNA methylation and gene expression data from various experiments. ARIC outperforms other methods through many evaluation metrics regardless of data types, especially for the estimation of rare component proportions. These results demonstrate ARIC as an effective tool to infer cell type fractions, especially in scenarios where rare components are of vital importance.

## Methods and Materials

### Problem definition

First, we modelled the deconvolution problem as a linear mixture of different cell types as previous studies did [20, 22]. Here we used ***m*** ∈ ℝ^*N*×1^ and 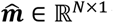 to denote the ideal value which do not contain any noise and the observed value of the bulk data, where *N* represents the number of markers. In order to estimate the proportion of cell types, we collected external references of possible components denoted as 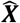, which is an *N* × *K* matrix where *K* is the number of cell types. The measured level of the *i*-th marker from the *j*-th cell type is denoted as 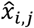 in the reference matrix 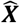. Similarly, the ideal external reference is denoted as ***X***. The ideal proportion of different cell types in one sample is denoted as a non-negative vector ***p*** ∈ ℝ^*K*×1^, where the sum of ***p*** should be equal to 1. We use 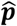 to denote the predicted value of ***p***. Under the assumption of linear mixing model, ***m*** can be regarded as the linear mixture of ***X***:

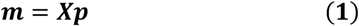

The task here is to estimate 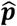 using 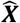 and 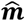.

### Employing component-wise condition number in eliminating feature collinearity

Solving ***p*** by Eq. 1 is an overdetermined equation problem due to the high-dimensional features of high-throughput omics data. Therefore, we first applied preliminary marker selection approaches to weaken the influence of biological and technical noise (see Supplementary Section 2). Even though, collinearity may occur in many features among different cell types, which may confuse the contributions of similar cell types and then lead to inaccurate proportion estimation. Previous studies [18, 29] used a condition-number-based strategy in RNA-seq data to measure the stability of the linear system against input variation or noise while reduce collinearity. As the measured ***m*** from a bulk sample is influenced by technical and biological noise, δ***p*** is defined as the change of proportion vector ***p*** if there occurs small changes or perturbations in the elements of ***m***, denoted as δ***m***. Besides, the relative errors of ***p*** and ***m*** are denoted as *Δp* and Δ*m* respectively. Then the condition number can be denoted as *C*:

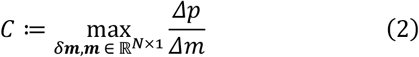

where

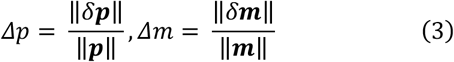

The condition number *C* measures the potential sensitivity of the proportion vector ***p*** to the change of bulk data ***m*** [30]. In particular, *C* takes the total relative error of the vector ***p*** into consideration, ignoring possible large relative errors on small-proportion components (see Supplementary Section 3).

In order to suppress the relative error of each component, here we used the component-wise condition numbers [31] to improve the procedure of marker selection. The definition of the component-wise condition number of the *c*-th component is:

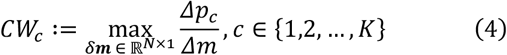

where

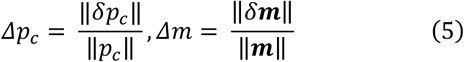

Different from Eq. 2, the numerator in Eq. 4 changes from Δ*p* to Δ*p* _*c*_, where Δ*p* _*c*_ denotes the relative error of component *c* in the proportion vector ***p***.

However, directly computing *CW*_*c*_ in Eq. 4 is impractical since it is a theoretical deduction with unknown content Δ*p*_*c*_. Therefore, we used Eq. 6 below to calculate *CW*_*c*_:

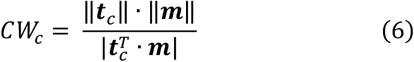

where 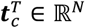 refers to the *c*-th row of ***X***^†^ which denotes the pseudo-inverse of ***X*** [31]. We computing *CW*_*c*_ using 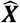 and 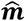 in practice.

We further calculated the largest condition number among all *K* components as the upper bound of relative errors, denoted as *CW*_*u*_:

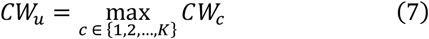

Since every component has a component-wise condition number, *CW*_*u*_ takes the most sensitive component into account. Through *CW*_*u*_, we were able to select markers avoiding large relative errors in any component. One example in Supplementary Section 4 illustrates how component-wise condition numbers successfully represent the large relative error on small components.

We eliminated markers leading to strong collinearity and used component-wise condition numbers to avoid large relative errors on each component. This process includes three main steps: (i) inspired by a former study [20], we used Euclidean distances to find cell type pair with the strongest collinearity. (ii) Next, the most similar marker of the cell type pair was eliminated and *CW*_*u*_ was computed. We used Eq. 8 to find the most similar marker *i*_*M*_ in cell type pair *j* and *k*:

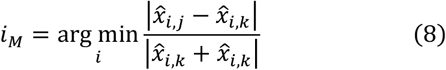

(iii)The above two steps were repeated to find the markers corresponding to the minimum *CW*_*u*_ . It should be noted that searching the global minimum *CW*_*u*_ is impractical, therefore we adopted a heuristic procedure to tackle this problem (see Supplementary Section 8).

### Adaptive and robust outlier detection

Through the above-mentioned marker selection procedure, we can select well-conditioned, non-collinear markers for each cell type. However, outliers that deviate from other markers may still exist, which may bring negative effects for precise deconvolution. Previously, few methods have tried to leverage outlier detection to improve the deconvolution performance. Here, we proposed a novel iterative and adaptive outlier detection method to overcome this problem based on robust regression [6, 32, 33].

We calculated the standardized residuals to indicate marker outliers. The standardized residual on the *i*-th marker is denoted as *SR*_*i*_, which can be written as:

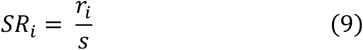

where *r*_*i*_ is the prediction error for bulk data, denoted as:

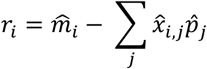

where 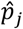 is the prediction value of the *j*-th component’s fraction. In addition, *s* is defined as:

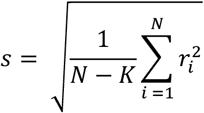

Then a decision whether marker *i* should be regarded as an outlier is made:

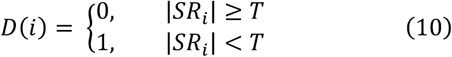

The threshold *T* was used to decide which marker should be treated as outliers and was a fixed parameter determined at the start of the outlier-removal approach (see Supplementary Section 8). Next, markers detected as outliers were removed and standardized residuals were recalculated based on remaining markers. This approach was iterated for ten times with the fixed *T*, and the rest markers were used for proportion estimation.

### Novel weighted SVR

Currently, υ-SVR has been proved to be a robust estimation method in many works [18, 34]. The primary problem of υ-SVR with linear kernel is as follows [35]:

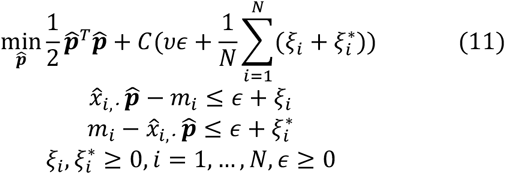

Here, 0 ≤ *v* ≤ 1, *C* is the regularization parameter. 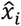, denotes the *i*-th row of 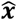, which represents the derived reference for marker *i*. The ε-insensitive loss function means that the loss is only considered when 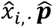 is beyond the range of *m*_*i*_ ± ϵ . Note that we use *m*_*i*_ instead of 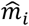 to denote the bulk data because 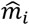 can be substituted by *m*_*i*_ after the previous feature selection.

From the Eq. 11, we can find that the absolute error term 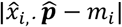 not only determines which markers should be the support vectors but also influence the loss function value. Inspired by dampened weighted least squares (DWLS) which adjust the weights of markers in the absolute errors loss function [36], we proposed a novel deconvolution method integrating marker weights with υ-SVR to minimize the relative errors on each component and avoid the ignorance of relative errors on rare cell types.

The absolute error term can be written as:

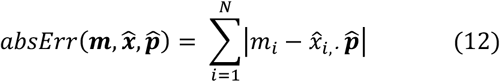

Same as DWLS, we define 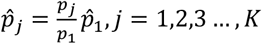. Then, rewrite Eq. 12:

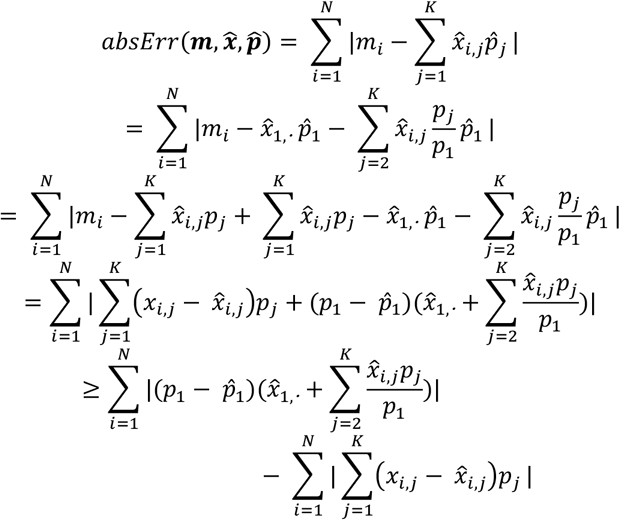

It is obvious that the second term is constant and the absolute error term is trying to minimize the errors on the first term. In consequence, we focus on the first term:

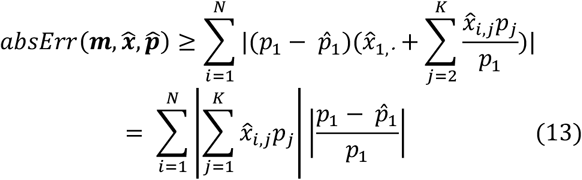

From Eq. 13, cell types which have larger reference value or a greater proportion lead to a larger impact on the absolute error term in υ-SVR and this phenomenon will lead to a biased estimation undoubtedly. Thus we designed weights for the markers to alleviate this problem:

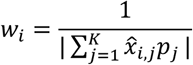

Then modify the absolute error term:

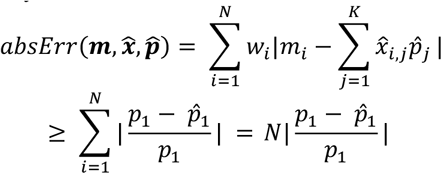

Without the loss of generality, we have the following relationships

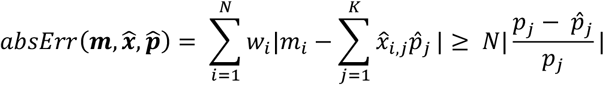

After weights adjustment, the absolute error term in υ-SVR can optimize the relative errors component-wisely, without ignoring rare cell types.

Finally, we normalize all the weights with Eq. 14 and apply υ-SVR to estimate proportions.

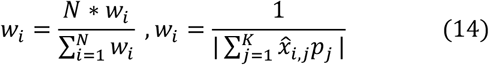

### Evaluation Metrics

Here we used the root mean square error (RMSE) and the Pearson’s correlation coefficient (PCC) as evaluation metrics as many former studies did [3, 18, 19]. Besides, in order to demonstrate the performance on different components, especially for rare components, we also adopted the mean absolute percentage error (MAPE) as another metric. The definition of RMSE and MAPE is:

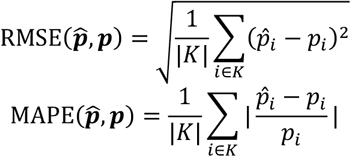

where 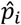 and *p*_*i*_ denotes the estimated value and the ground truth of each component respectively.

### Benchmark methods

We selected state-of-art deconvolution methods that were designed for DNA methylation data or gene expression data for benchmark. Benchmarks for analyzing DNA methylation data include QP (quadratic programming) [37], Moss [20], Epidish (robust partial correlation) [38, 39], Sun [21], MethylCIBERSORT [40] and MethylResolver [41]. Benchmarks for analyzing gene expression data include QP [37], EPIC [19], CIBERSORT [18], dtangle [3], FARDEEP [6] and DeconRNASeq [42]. Of course, some methods which are already compared with other methods in previous works [19, 43] are not included in our analysis. Different preliminary marker selection strategies were used in different kinds of data (see Supplementary Section 2).

### Datasets

We selected state-of-art HumanMethylationEPIC BeadArray data were used in both simulation and real data evaluation [44]. Data used for *in silico* simulation contains six cell types: NK cells, Neutrophils, B cells, monocytes, CD4^+^ T cells and CD8^+^ T cells, each cell type with six samples. Additionally, there are 12 real samples which are the mixtures of genomic DNA from six purified leukocyte subtypes, with known proportions for evaluation [44].

153 paired-end RNA-seq samples of 8 cell types were collected (Supplementary Table 2), including B cells, CD4^+^ T cells, CD8^+^ T cells, endothelial cells, macrophages, monocytes, neutrophils and NK cells. We processed raw RNA-seq data into transcripts per million (TPM) to generate simulation data [19]. Single-cell RNA-seq (scRNA-seq) data were adopted from a previous study [45]. We selected scRNA-seq data of 7 tissues from 6 to 10-week-old female mice for generating pseudo-bulk data.

Four gene expression microarray datasets and three RNA-seq datasets with known proportion are collected.

All the datasets can be accessed through supplementary table 1∼4.

## Results

### ARIC infers the cell type fraction accurately on in silico mixed data

We first evaluated ARIC on different *in silico* mixed datasets including DNA methylation assays, RNA-seq and scRNA-seq. Each dataset was divided into two parts, one for constructing the external reference and the other for evaluation. Proportions of all components were generated randomly and adequate samples were produced to ensure the existence of rare components (see Supplementary Section 7). RMSE, MAPE and PCC were calculated to evaluate the performance of different methods (Fig. 1).

**Fig. 1.**
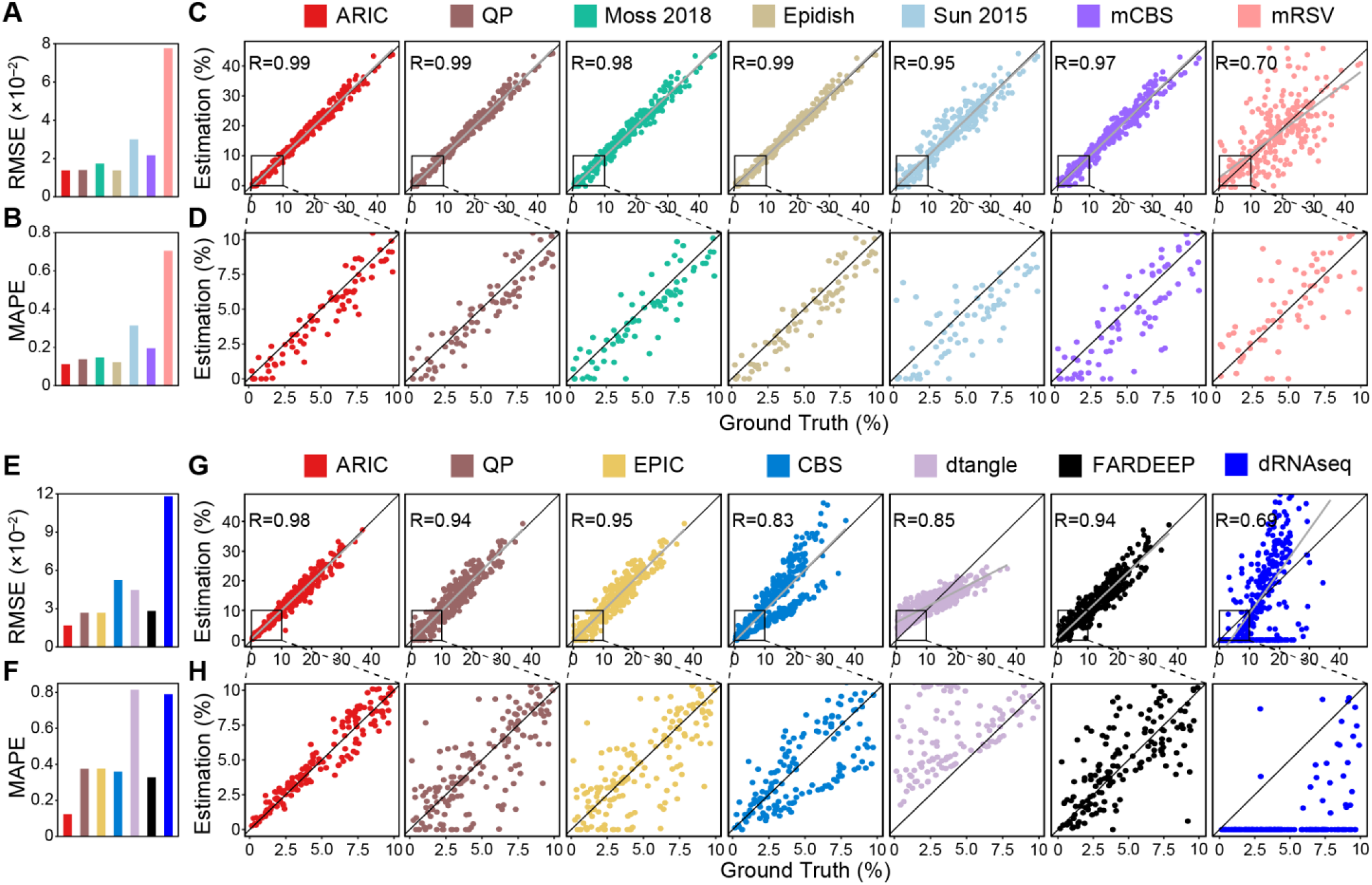
Deconvolution results for *in silico* bulk DNA methylation (A∼D) and gene expression (E∼H) datasets. (A) and (E) the RMSE for each method. (B) and (F) the MAPE for each method. (C) and (G) are the scatter plots for estimated cell-type fractions against true fractions. PCCs are shown in the top-left corner of each panel. (D) and (H) are zoomed-in versions of (C) and (G) for cell-types with fraction less than 10%. Gray lines represent linear regressions of the points. mCBS: MethylCIBERSORT, mRSV: MethylResolver, CBS: CIBERSORT, dRNAseq: DeconRNASeq.

Though PCCs of some methods are comparable with ARIC, ARIC achieves a relatively low RMSE and MAPE compared with any other methods for both DNA methylation assays (Fig. 1A-B) and RNA-seq (Fig. 1E-F), which indicates the high accuracy of ARIC for the prediction of every component’s fraction. What’s more, as shown in Fig. 1C∼D and G∼H, the prediction results of ARIC are squeezed into the diagonal line gradually as the true fraction decreased by contrast with other methods, which indicates the precision of ARIC on rare components. Interestingly, the prediction results of dtangle showed a significant deviation in comparison with the ground truth (the last panel in Fig. 1G), though with a high PCC.

To further evaluate the performance of ARIC, we generated pseudo-bulk RNA-seq samples *in silico* with data adopted from a well-characterized scRNA-seq study [45]. As shown in Supplementary Fig. S1, ARIC outperforms in metrics compared with others, especially for rare components. Interestingly, the deviation of dtangle’s prediction still exists. dtangle shows a higher PCC as well as a smaller RMSE than all the other methods except for ARIC and FARDEEP, but exhibits the highest MAPE value in the meantime. The results, together with the results shown in Fig. 1G, indicate that PCC and RMSE are insufficient to depict the prediction accuracy and may cause erroneous judgements.

### ARIC estimate rare cell type fractions accurately

To illustrate the capability of ARIC to deal with rare components, we computationally mixed methylation data [20] of individual cell types at varying proportions. *In silico* simulations were performed by setting the proportion of one component to 1%, 3%, 5%, 7% and 10% respectively, and the proportions of other components were generated randomly. We evaluated the results of the rare component solely with all the three metrics (Fig. 2). ARIC always shows the smallest RMSE (Fig. 2A) and MAPE (Fig. 2B) among all methods across different rare-component proportions. Moreover, the results of ARIC are the closest to the ground truth with the smallest variance across different proportions (Fig. 2C), which illustrates the robustness of ARIC on inferring the fraction of rare components. The same results were obtained when inferring components with lower proportions from 0.1% to 1% (Supplementary Fig. S2).

**Fig. 2.**
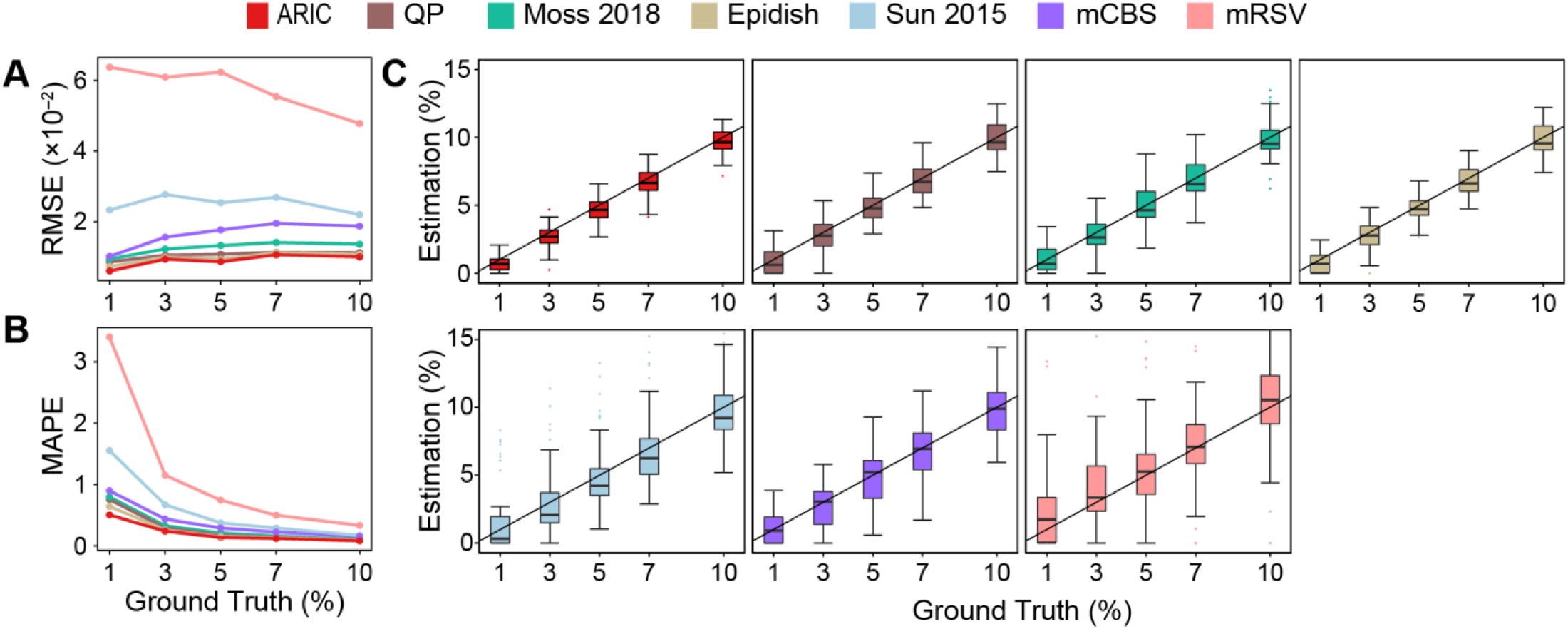
Deconvolution results for simulated rare components varying from 1% to 10%. (A) and (B) show RMSEs and MAPEs gradually changing with the rare component proportion. (C) is the box-plot of the deconvolution results with replicates (*n* = 60). Black lines represent the estimation values are equal to the ground truth. mCBS: MethylCIBERSORT, mRSV: MethylResolver.

### ARIC outperforms in the deconvolution of real data from multiple sources

To evaluate the efficacy of ARIC on real data, we collected a DNA methylation dataset [44], which provides us 12 samples with known fractions of each cell type. The deconvolution results are shown in Fig. 3. Though all methods exhibit high PCCs, ARIC shows a better performance than any other baseline methods on the metrics of both RMSE and MAPE.

**Fig. 3.**
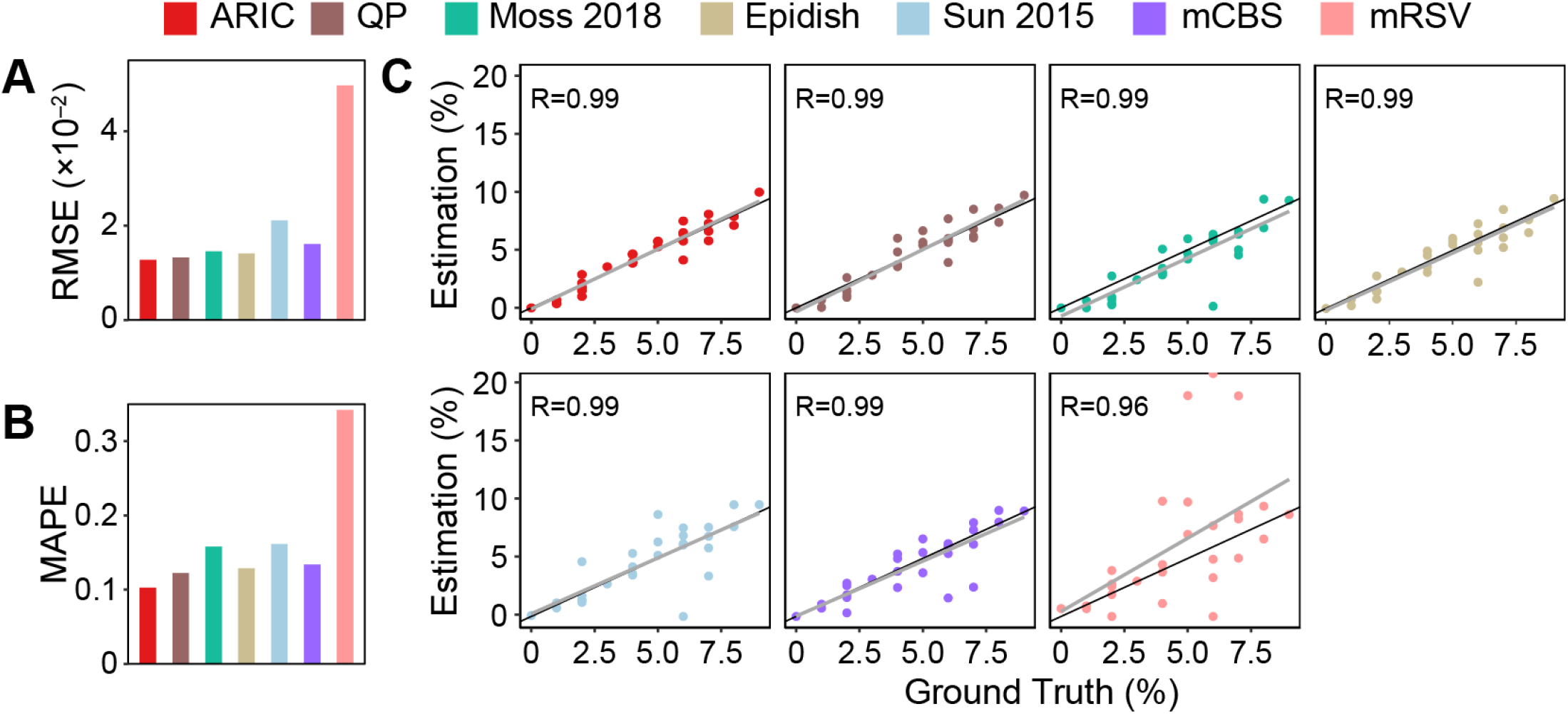
Deconvolution results for experimentally mixed methylation data. (A∼C) show RMSEs, MAPEs and prediction results against the ground truth respectively. PCCs are shown in the top-left corner for each method in (C). Gray lines in (C) represent linear regressions of the points. Black lines in (C) represent the estimation values are equal to the ground truth. mCBS: MethylCIBERSORT, mRSV: MethylResolver.

We also collected seven expression datasets measured by microarray or RNA-seq with known cell type fractions. We calculated the performance of each method on all datasets and analyzed the results jointly. ARIC achieves a relatively lower RMSE (Fig. 4A) and MAPE (Fig. 4B) in different datasets. The median RMSE and MAPE of dtangle are almost the same as ARIC, but our method shows more outstanding outcomes when focusing on rare components (Fig.4C-D). ARIC also shows the smallest deviation among all methods (Fig. 4), which indicates the robustness of ARIC.

**Fig. 4.**
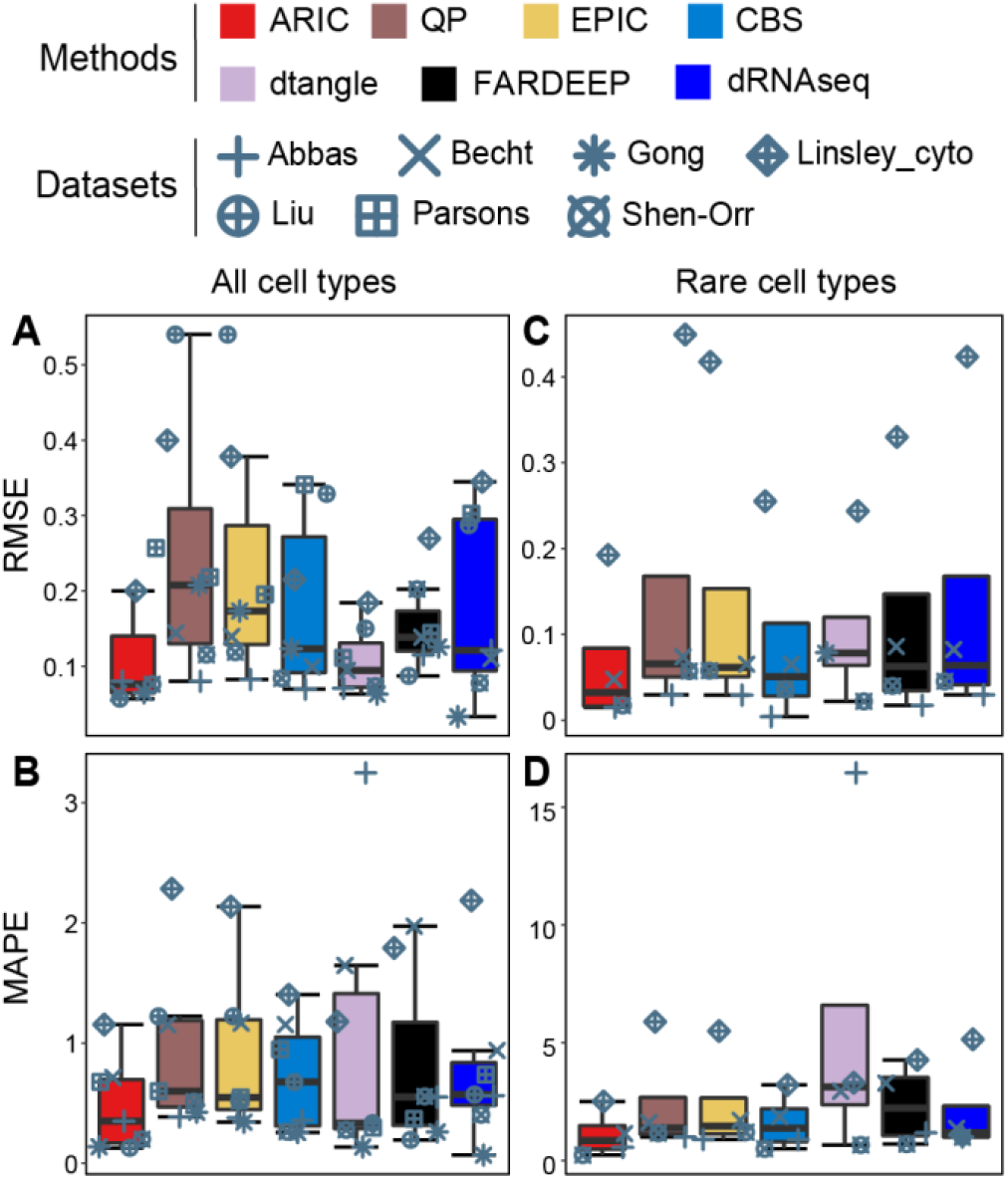
Deconvolution results for experimentally mixed expression data. (A) and (B) are RMSEs and MAPEs for all components. (C) and (D) are RMSEs and MAPEs for rare components with fractions less than 10%. Black lines represent the median for each method. CBS: CIBERSORT. dRNAseq: DeconRNASeq.

## Acknowledgements

We would like to thank Dr. Qiongye Dong and Ms. Jiaqi Li for helpful discussions.

## Discussion

As a new approach for deconvolution of cell type fractions, ARIC is tested on both *in silico* mixed data as well as real data with variou0073 DNA methylation and gene expression datasets. Several benchmarks are selected to compare with ARIC. For all kinds of datasets, ARIC shows the ability to estimate the fractions of different cell types, especially for rare components, more precisely than any other deconvolution methods.

The remarkable performance of ARIC owes to several novel designs in the algorithm. Different cell types that are differentiated from same progenitors or share similar functions may exhibit similar methylome or transcriptome profiles, which may further lead to confounded deconvolution results due to collinearity [36]. Previous studies minimized the condition number to improve the accuracy of the deconvolution results [18, 34]. However, the condition number cannot pay equal attention to all cell types as it measures the overall error of all components instead of the component-wise error. This will cause a strong bias for rare cell types. Therefore, the component-wise condition number is introduced into ARIC to evaluate the error of each component more equally to get rid of the negligence of rare components. During the marker selection process, the component-wise condition number is calculated at each step, and then markers leading to the smallest component-wise condition number are adopted for further analysis. The deconvolution results of both simulated and real datasets reveal that employing the component-wise condition number brings ARIC a more powerful capacity to estimate the fraction of rare components.

Furthermore, robustness is an indispensable requirement for bulk data deconvolution to ensure high accuracies. However, the deconvolution procedure is susceptible to outliers brought by measure error or environmental effect, which may hamper the robustness of algorithms [6]. Here, ARIC utilizes the standardized residual to distinguish outliers from effective markers. Notably, some outliers are hard to be differentiated when there exist more significant outliers in all markers. As the consequence, outliers can hardly be detected and removed without iterations. Therefore, ARIC computes standardized residuals and detects outliers adaptively to ensure that outliers are removed as precisely as possible.

There is still room for improvement of ARIC. ARIC depends on external references whose quality may influence the prediction results significantly. As the rapid development of single-cell sequencing technologies, purer references can be produced to enhance the performance of ARIC. Moreover, as ARIC is developed independent from the type of data and performs well in data generated from HumanMethylationEPIC BeadArray, microarray, RNA-seq and scRNA-seq, ARIC is promising to be applied into other bulk data, such as ATAC-seq and FAIRE-seq.

In conclusion, ARIC is a robust and accurate tool for decoupling the fraction of cell types in mixture data. Particularly, ARIC can estimate the fraction of rare components far more precisely and robustly than other methods, suggesting ARIC as a promising tool to solve the deconvolution problem of bulk data where rare components matter, which may further benefit both scientific research and clinical applications.

## Notes

### Competing Interest Statement

The authors have declared no competing interest.

