## Supplementary for "ARIC: Accurate and robust inference of cell type proportions from bulk gene expression or DNA methylation data": Supplementary.pdf

#### 1 Datasets Collection and Processing

##### 1.1 MethylationEPIC BeadChip Data

All the EPIC chip data [1] are download from NCBI, samples we used can be found in Supplementary table.1.

We use R package ChAMP [2] with default parameters to do all the pre-processing and extract methylation level for every single CpG site.

##### 1.2 Gene Expression Microarray

All the microarray data [3-6] are downloaded from GEO and samples we used can be found in Supplementary table.5.

##### 1.3 RNA-seq and scRNA-seq

8 cell type related RNA-seq samples are carefully collected, all samples can be found in supplementary table.2 [7-16]. We collected every cell type at least from two different experiments or laboratory in order to get rid of overfitting from the same experiment. We only use paired data to avoid influence raised from different data type like single and paired sequencing. Fastqc [17] are used for quality control. Low quality data and low depth data will be discarded. Adapters are detected and removed using Adapterremoval [18]. Clean read are aligned using Hisat2[19] with parameter "-q --no-mixed --no-discordant" and read are counted by featureCounts [20] with annotation file Homo_sapiens.GRCh37.75.gtf. Differential expression analysis are performed using DEseq2 [21] with default parameters. Read counts are converted to Transcripts Per Million (TPM) using customized script as the former work [22].

3 known proportion RNA-seq studies [16, 23, 24] are used to evaluate the deconvolution performance. All samples can be found in supplementary table.2.

Single cell RNA-seq data are from the previous work [25]. We selected 7 cell types (see supplementary table.3) from same sourcement. We use the raw counts data provided by the author in our analysis.

#### 2 Preliminary Marker Selection

Due to different properties of data we used, we apply different marker selection strategies.

##### 2.1 MethylationEPIC BeadChip Data

Due to different properties of data we used, this part can be divided into two categories: methylation data and gene expression data.

In our research, we use methylation chip data to evaluate our algorithm, so we decide to use ChAMP package [2] in R language to assist us with data pre-processing. The main step here is to perform filtering according to p-value and the number of beads of probes. Normalization is also necessary for probes in different design types. After pre-processing, a one-versus-one marker selection strategy is developed to perform preliminary marker selection, which includes two steps. First, we define every two cell types as one pair. Second, for each pair, we compute variation to find top two hundred different markers.

$$V_{i,j}^{s} = \frac{x_{s,i}}{x_{s,i} + x_{s,j}}$$

where $V_{i,j}^{s}$ denotes variation, i and j refers to the index of cell types, s denotes the s'th marker and $x_{s,j}$ means the value of the reference in the $i$-th row and $j$-th column.

We define top 200 markers as the markers with highest and lowest 100 $V_{i,j}^{s}$ respectively. Due to the symmetry of $V_{i,j}^{s}$, the order of i and j does not matter for preliminary selection. Then this step should be finished by incorporating all markers from every pair into one set.

#### 2.2 RNA-Seq simulation data

For RNA-Seq data we directly leverage methods in EPIC [22] to perform preliminary marker selection. These marker genes are specific to cell types, which can reduce the collinearity for every method as much as possible. To make sure we have a reliable evaluation, all methods take the same marker genes as input.

#### 2.3 scRNA-Seq simulation data

We ranked the differential expressed genes of scRNA-seq data according to differential expressed p-value across cell types and select the top 500 genes as input for all evaluation methods.

#### 2.4 Other datasets

We also perform evaluation on other datasets, like real gene expression data. For these datasets we select markers according to the expression variance between different cell types (differential expressed gene of different cell types). The most variable markers are selected as input. In our work, we selected top 1000 markers.

### 3 Small Proportion Problem in Deconvolution

Previously many works chose Root Mean Square Error (RMSE) as their metric which only reflect the error of the whole vector regardless of each component within the vector. This may result in such situation: the estimated whole proportion vector has a small relative error, but for some small components, the departure may be several times from the ground truth. We provide an example here, $p$ and $\hat{p}$ illustrate the ground truth and prediction respectively.

$$p = \begin{bmatrix} 0.01 \\ 0.2 \\ 0.5 \\ 0.29 \end{bmatrix} \quad \hat{p}_1 = \begin{bmatrix} 0.05 \\ 0.16 \\ 0.5 \\ 0.29 \end{bmatrix} \quad \hat{p}_2 = \begin{bmatrix} 0.01 \\ 0.24 \\ 0.46 \\ 0.29 \end{bmatrix}$$

$\hat{p}_1$ and $\hat{p}_2$ are the two different prediction. Now let's look at RMSE; for $\hat{p}_1$, RMSE = 0.0283, for $\hat{p}_2$, RMSE = 0.0283; so we cannot see any difference just from RMSE, however, if we check every component, we can find the first component of $\hat{p}_1$ has a much larger relative error than $\hat{p}_2$; this phenomenon can be observed by looking at the Mean Average Percentage Error (MAPE); for $\hat{p}_1$, MAPE = 1.05, for $\hat{p}_2$, MAPE = 0.07.

### 4 Example of Component-wise Condition Number

One example in Supplementary illustrates how the component-wise condition number successfully captures the potential large relative error on small components. We generated simulated reference $R_s$ and simulated mixture data $M_s$:

$$R_s = \begin{bmatrix} 0.9 & 0.3 & 0.8 & 0.3 \\ 0.1 & 0.2 & 0.5 & 0.9 \\ 0.4 & 0.3 & 0.8 & 0.1 \\ 0.7 & 0.8 & 0.7 & 0.9 \end{bmatrix} \quad M_s = \begin{bmatrix} 0.6495 \\ 0.1804 \\ 0.3933 \\ 0.7402 \end{bmatrix}$$

We assign the proportion data $P_s = [0.5, 0.4, 0.099, 0.001]$, where the 4th component can be seen as the tiny component. Perturbations are added to $M_s$. By solving the equation, we can compute the relative error: $\Delta p = 0.051$ and traditional condition number: $C = 8.6226$. It seems that perturbations have little influence on results. However, if we look into every component:

$$\begin{bmatrix} \Delta p_1 \\ \Delta p_2 \\ \Delta p_3 \\ \Delta p_4 \end{bmatrix} = \begin{bmatrix} 0.0207 \\ 0.0562 \\ 0.1363 \\ 20.8703 \end{bmatrix}, \begin{bmatrix} CW_1 \\ CW_2 \\ CW_3 \\ CW_4 \end{bmatrix} = \begin{bmatrix} 5.3486 \\ 8.3605 \\ 22.4224 \\ 1706.2 \end{bmatrix}$$

This example shows how the component-wise condition number successfully captures the potential larger relative error on small components. With the assistance of component-wise condition number, we are able to take all components into consideration by picking the largest condition number among all components as the upper bound. $CW_u$ denotes the largest componentwise condition number.

### 5 Support Vector Regression Parameters in RAIP

When outliers have been removed, we apply weighted nuSVR in our deconvolution process from python package sklearn. We set the parameter nu = 0.25, 0.5 and 0.75 respectively.

### 6 Benchmark Methods

We have known different deconvolution scenarios have different methods, in order to demonstrate capability of RAIP over other methods, we chose different baselines for methylation data and gene expression data.

For methylation data, we select QP (quadratic programming) [26], Moss [27], Epidish (robust partial correlation) [28, 29], Sun [30], MethylCIBERSORT [31] and MethylResolver [32] as benchmark methods; QP and Epidish can take the markers we select as input, so we use the same markers as RAIP for these methods; however, Moss, Magenheim [27] and Sun, Jiang [30] cannot be provided with our markers because they have their own rules for markers selection.

MethylCIBERSORT and MethylResolver are executed in default mode.

For gene expression data, including RNA-Seq data, scRNA-Seq data and gene expression microarray data, we compare RAIP with QP [26], EPIC [22], CIBERSORT [33], dtangle [34], FARDEEP [35] and DeconRNASeq [36], which has been widely used and proved to be effective. All of these markers can be provided with markers we select, so we keep these methods with the same markers.

### 7 *In Silico* Simulation

For all simulation, we divided our dataset into two parts: training set and test set. Training set is used to construct the external reference panel and test set is responsible for evaluation by performing simulated mixing.

#### 7.1 MethylationEPIC BeadChip Data

Methylation chip data our used contains 6 sample for every cell type. Therefore, we use 4 samples of every cell type to build external reference and 2 samples to generate mixture data. The mixture proportion are generate from uniform distribution U(0, 1000). For a fixed reference, 50 different mixture proportion samples are generated.

#### 7.2 RNA-seq and scRNA-seq Data

We separate every cell type RNA-seq samples into train (70%) and test (30%). For train set, we compute average TPM value for every gene as external reference penal. Test set are used for generate mixture data. We believe that this method remains individual variance as well as biological noise. The mixture proportion are generate from uniform distribution U(0, 1000). For a fixed reference, 50 different mixture proportion samples are generated.

scRNA-seq simulation data were generated in the same way.

#### 7.3 Small proportion component simulation

We leverage MethylationEPIC BeadChip Data to construct the small proportion simulation data. In this part we set one cell type's proportion to the small proportion value, and other cell types' proportion were generated from uniform distribution. This procedure was performed for each cell type, 10 samples were generated once, 60 samples together.

We set small proportion values to 0.001, 0.003, 0.005, 0.007, 0.01 separately. Therefore, we get 300 samples totally.

### 8 RAIP Related Parameters

For first marker selection step (main text section "Employing component-wise condition number in eliminating feature collinearity"), it is notable that there is a trade-off of selecting repeat times: we should consider both the collinearity factor elimination and adequate markers containing enough information for fractions inference. W set 10% of these marker number as the iterative number during integrating component-wise condition number into marker selection procedure.

For second marker selection step (main text section "Adaptive and robust outlier detection"), we set threshold $T$ to the value which can remove 25% markers in the first loop. Actually, $T$ should be set to 2.5 as the previous work described [37]. However, in our massive testing, setting $T$ based on the number of markers removed in the first loop can be well adapted to different datasets.

To avoid some of the weights too large in υ-SVR, we set a threshold in practice:

$$w_i = \begin{cases} \frac{1}{|\sum_{j=1}^{K} \hat{x}_{i,j} p_j|}, & if \quad \frac{1}{|\sum_{j=1}^{K} \hat{x}_{i,j} p_j|} < 10 * \min(w_1, \dots, w_N) \\ 10 * \min(w_1, \dots, w_N), & else \end{cases}$$

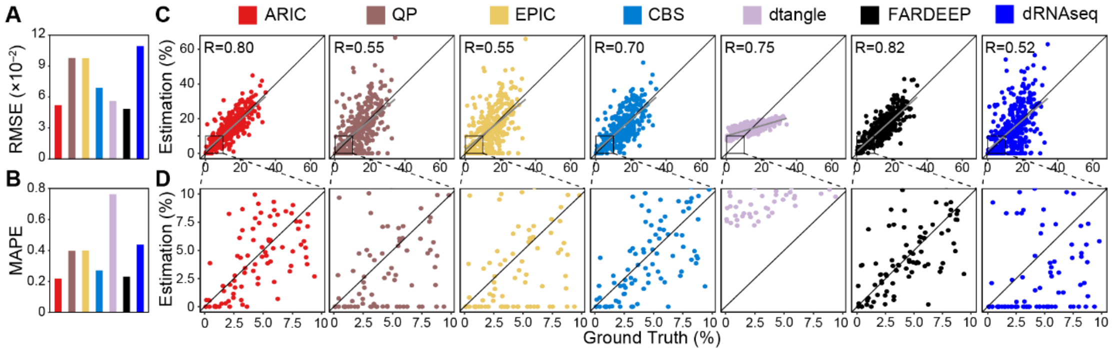


**Fig. S1** Deconvolution results for scRNA-seq data. (A) is the RMSE for each method. (B) is the MAPE for each method. (C) the scatter plots for estimated cell-type fractions against true fractions. PCCs are shown in the top-left corner of each panel. (D) is zoomed-in versions of (C) for cell-types with fraction less than 10%.CBS: CIBERSORT. dRNAseq: DeconRNASeq.

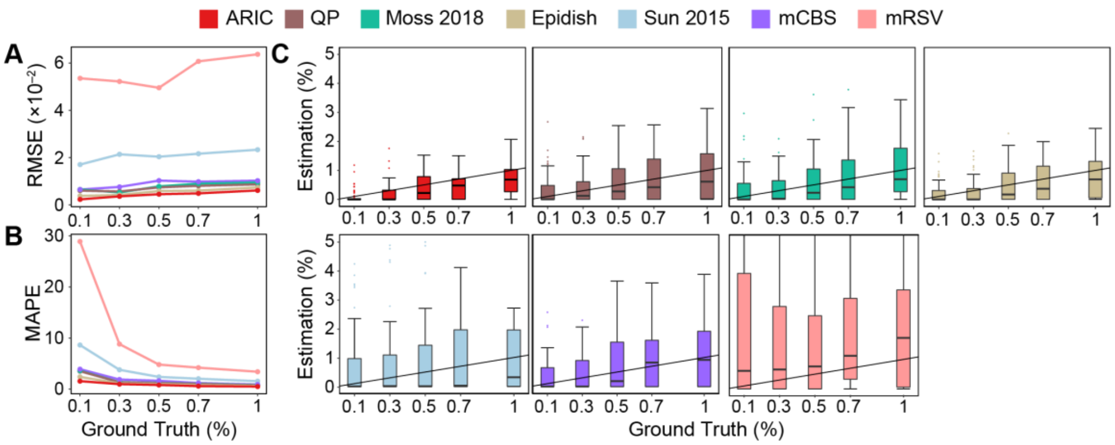


**Fig. S2** Deconvolution results for simulated rare components varying from 0.1% to 1%. (A) and (B) show RMSEs and MAPEs gradually changing with the rare component proportion. (C) is the box-plot of the deconvolution results with replicates (n = 60). Black lines represent the estimation values are equal to the ground truth. mCBS: MethylCIBERSORT, mRSV: MethylResolver.
